# F-actin dynamics transform filopodial bridges into intercellular nanotubes capable of distant cell communication

**DOI:** 10.1101/405340

**Authors:** Minhyeok Chang, Jaeho Oh, Junsang Doh, Jong-Bong Lee

## Abstract

A novel actin-based bridge connecting cells has been recognized as a new pathway for the distant transport of cytoplasmic components, viruses, or pathogenic substances between cells. However, it is not yet known how such a fine structure extends over several hundred micrometres and remains robust for several hours. Using optical fluorescence imaging methods, we found that random contact promotes the formation of filopodial bridges through N-cadherin interactions between filopodia, which are slender actin-rich plasma membrane protrusions. These filopodial bridges eventually evolve into a single actin-based bridge (intercellular nanotube) that connects two cells via an intermediate state that involves a helical structure. Surprisingly, the twisting of two filopodia is likely to result from the rotational motion of actin filaments inside the filopodia by myosin V. The accumulated torsion of the filopodia triggers the release of one of the paired filopodia, whose end is attached to the other cell body by an N-cadherin cluster. The resulting retraction of the filopodium by retrograde F-actin flow leaves a single bridge. The N-cadherin/catenin cluster is likely to form a synapse between the intercellular nanotube and the cell body. This study sheds light on the formation mechanism of the filopodial bridge-based intercellular nanotubes for long-distance communication between cells.

## Introduction

In recent decades, intercellular bridges that connect cells over long distances of up to several cell diameters have been intensively investigated to verify their biological significance. Receptors^1-5^, organelles^6, 7^, vesicles^8, 9, viruses^10, 11^, morphogens^12-15^, prions^16^ and nucleic acids^17-19^ can be transported through these unique structures in various types of cells, including neuronal cells^6, 7, 20^, immune cells^8, 11, 21^, developing cells^12-15^, cancer cells^17, 22, 23^, epithelial cells^24^ and live *Drosophila* tissue^1-5, 18^. Moreover, a recent study on primary cancer cells showed the medical importance of intercellular nanotube (INT) formation in tumour metastasis^17^. These actin or microtubule bundle-based intercellular structures have been named tunnelling nanotubes (TNTs)^6, 7^, 23-27^, TNT-like structures^12^, membrane nanotubes^8, 11, 21, 22, 28^, nanotubes^15^, filopodial bridges (FBs)^10^ or cytonemes^1-5^, depending on the topology. INTs have been reported as a specific and distant route for intercellular communication, in contrast to long-distance but nonspecific (soluble factors or exosome) or specific but short-distance (various types of intercellular junctions or synapses) interactions between cells^29-31^.

Imaging studies on cultured cells suggested that INTs can initially form from the interaction between thin finger-like actin assembly-driven protrusions (filopodia) or from direct contact between plasma membranes^20, 28^. The withdrawal of the cells from each other eventually generates long and linear bridges. The resulting INT is suspended between the cells^6^ while maintaining a length a few hundred times longer than its thickness. Since filopodia exhibit various physical features, such as growth^32^, retraction^33^, bending^34^, rotation^35^ and helical buckling^36^, it would be rational to assume that the formation of INTs is most likely to be dynamic. These physical properties and the formation mechanism of INTs have rarely been investigated, as most INT studies have focused on the biological roles of INTs in intercellular communication.

We visualized the process of INT formation in HeLa cells and identified the structure of INTs and their configuration in an intermediate state using real-time fluorescence microscopy and super-resolution fluorescence microscopy^9, 28, 37^. Here, we denote an INT as a single protrusion connecting two cells. We found that the interaction of N-cadherin molecules on filopodium membranes maintains contact between two filopodia, which results in a filamentous actin (F-actin) bridge (filopodial bridge). We observed that the FB appears to be twisted around itself in an intermediate state, and this effect results from actin polymerization and the rotation of the actin filaments inside the filopodia by myosin motor proteins in lamellipodia. The torsional energy in the FB triggers the separation of the filopodia after one of the filopodia reaches and tightly binds to a cell body via an N-cadherin/β-catenin cluster. The retraction of the released filopodium ultimately allows the formation of an INT that connects the two cells. These imaging studies reveal the mechanism of INT formation and suggest an intercellular nanostructure that may play a critical role in the formation of long-standing INTs between cells.

## Results

### Dynamic transition of FBs between cells

To visualize the dynamic formation of actin-based INTs in living cells, we tagged cellular F-actin with an actin-binding 17-amino-acid peptide (Lifeact) that was genetically engineered to contain GFP (green) or mCherry (red)^38^ (Fig. 1a; Methods). Cells transfected with Lifeact-GFP or Lifeact-mCherry were cultured together at a 1:1 ratio (Methods). Actin-based protrusions from cells were imaged using line-scan confocal microscopy (LSCM; Supplementary Fig. 1; Methods)^39^. The colocalization images illustrate two-coloured bridges formed by two filopodia protruding from different cells (FBs, Fig. 1a top) or single-coloured bridges between cells (INTs, Fig. 1a bottom). FBs were observed more frequently than INTs in live or fixed cells after 12 hr of INT stimulation and 24 hr of co-culture (Methods), which is nearly identical to the frequency of FBs and INTs obtained by visualizing the plasma membrane after labeling with the DiD and DiI dyes (Supplementary Fig. 2). We confirmed from this plasma membrane imaging that a FB is not a single membrane tube formed by the membrane fusion of two filopodia but a bridge consisting of two isolated filopodia (Fig. 1a, left cartoon). Interestingly, the frequency of INTs significantly increased from 33 ± 6% to 50 ± 6% as the INT stimulation time increased from 12 hr to 36 hr (Fig. 1b; Methods).

**Fig 1.**
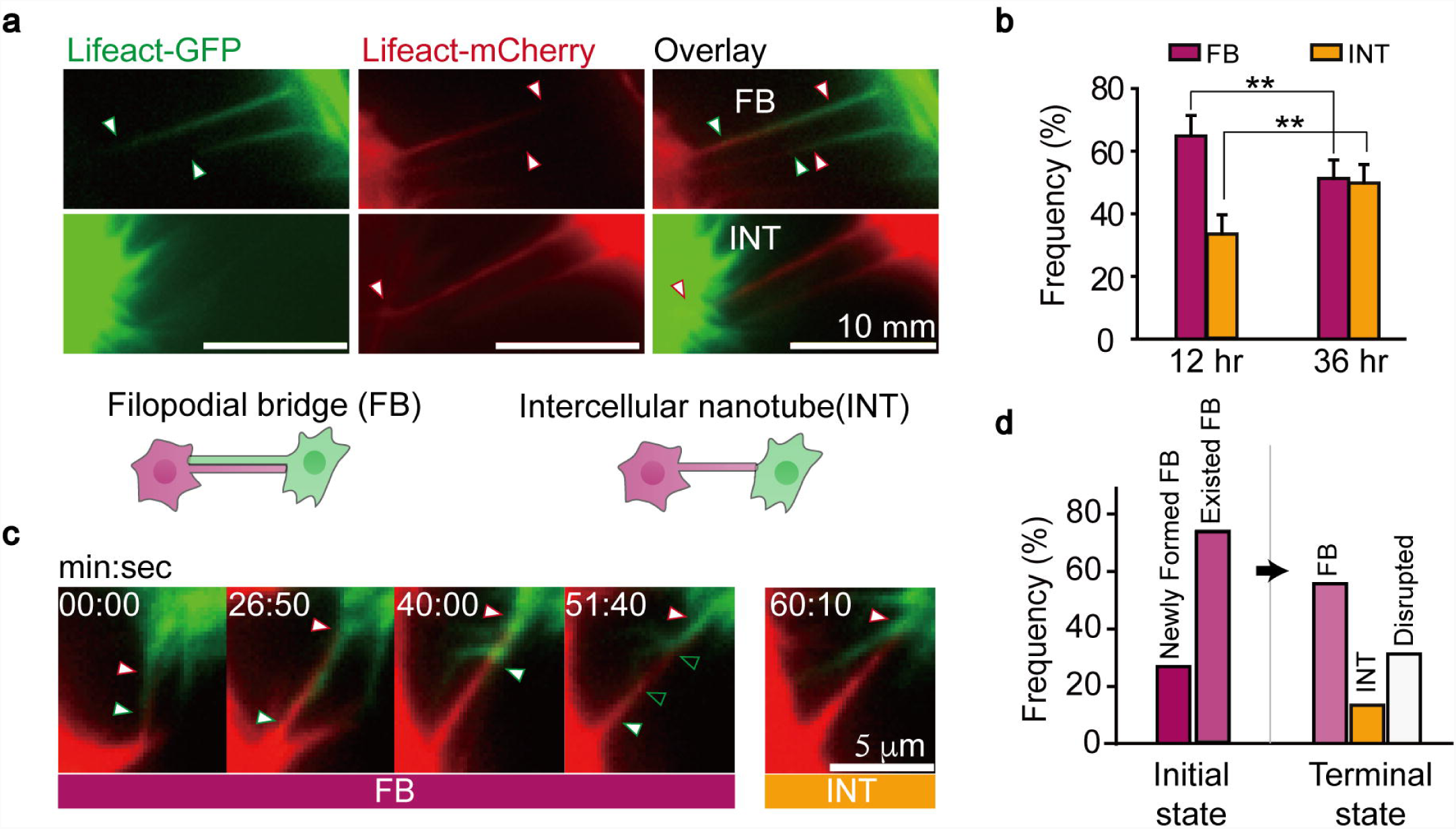
Dynamic transition from FBs to INTs. **a**, Colocalization of F-actin genetically engineered to contain Lifeact-GFP or Lifeact-mCherry in each HeLa cell. The triangles indicate the end of each actin protrusion (filopodium) that consists of intercellular connections. We denote that in contrast to the filopodial bridge (FB) formed by two filopodia protruding from each cell, the intercellular nanotube (INT) consists of a single membrane protrusion connecting two cells. **b**, Frequencies of FBs and INTs in cells fixed at different times (after 12 hr and 36 hr of INT stimulation 24 hr after cell seeding), showing a significant increase in the frequency of INTs from 33% to 50% (n = 242 and 158 INTs obtained from N_cell_ = 116 and 60 cell pairs in N_exp_ = 4 independent experiments). \*\**p*-value < 0.01 by Student’s t-test. The error bars indicate the s.d. **c**, Transition dynamics of FBs to INTs during real-time imaging of live HeLa cells. A filopodium retrogrades along the paired filopodium (40:00 and 51:40; green) and then a single bridge is eventually formed (60:10; red). In live cell imaging, the formation of an INT without an FB has never been observed (N_cell_ = 20). **d**, Some newly formed (n = 11) or pre-existed (n = 33) FBs developed into new INTs (n = 6, 13%) or were disrupted (n = 14, 31%), while others remained in the FB states (n = 25, 56%). The cells were imaged in 12 - 36 hr of INT stimulation 24 hr after cell seeding (N_cell_ = 17) for time-lapse experiments (recording every 5 or 10 s with a 100 ms exposure time).

Real-time tracking of membrane protrusions revealed the initial and intermediate states of INT formation (Fig. 1c). Contact between fluctuating filopodia protruding from green and red cells formed FBs (Fig. 1c, 00:00; Supplementary Movie 1). The resulting FBs appeared to extend until one or both of the filopodia reached the other cell body (Fig. 1c). One filopodium was then released from the other filopodium and retracted back to the cell body such that a single filopodium remained connected between the two cells to finally form an INT (Fig. 1c; Supplementary Fig. 3a). The contact and dislodgement of lamellipodia from two cells also initiated FB formation (Supplementary Fig. 3b; Supplementary Movie 2). However, we did not observe direct formation of an INT from the extension of a single filopodium or lamellipodium (Supplementary Fig. 3c). Furthermore, we observed that FBs developed into INTs or were disrupted, leaving individual filopodia, in time-lapse imaging of live cells for 1 hr after at least 12 hr of INT stimulation (Fig. 1d; Supplementary Fig. 3d; Supplementary Movie 3). Highly dynamic F-actin motion is likely to enhance the physical contact of filopodia for FB formation. In fact, breakage of FBs, which results in two separate filopodia, often occurred due to the large fluctuations of the filopodia (Fig. 1d, 31%; Supplementary Movie 1). Taken together, these results strongly suggest that INTs result from the transition of FBs formed by the dynamic motion of two filopodia.

### N-cadherin interactions between two filopodia support INT formation

Binding of the extracellular domains of cadherin molecules for cell-to-cell adhesion is a promising potential mechanism of FB formation and stabilization^40^. N-cadherin, which is a classical cadherin expressed in HeLa cells, and F-actin were colocalized in fixed cells labelled with an Alexa Fluor 488 (AF488)-conjugated antibody specific for N-cadherin and AF647-phalloidin for F-actin staining (Fig. 2a; Methods). Interestingly, N-cadherins were uniformly distributed along the bridge connecting two cells (Fig. 2a, top; Fig. 2b, 70%; ‘Spread’) or localized at the end of the bridge (Fig. 2a, bottom; Fig. 2b, 30%; ‘Cluster’). Cadherin molecules form a complex with a heterodimer of α-β-catenin that binds to F-actin for the cadherin-cadherin binding-mediated adhesion of plasma membranes^41^. Fig. 2c shows that the N-cadherin and β-catenin signals were strongly correlated on FBs or INTs [Pearson’s correlation coefficient = 0.54 ± 0.30; mean ± standard deviation (s.d.); Methods]. To quantify the degree of local colocalization of N-cadherin and β-catenin, we defined a local correlation coefficient at each pixel using the intensity profiles obtained from the colocalization image (Methods). The N-cadherin/β-catenin correlation appeared to be strong near the cell body (Fig. 2d).

**Fig 2.**
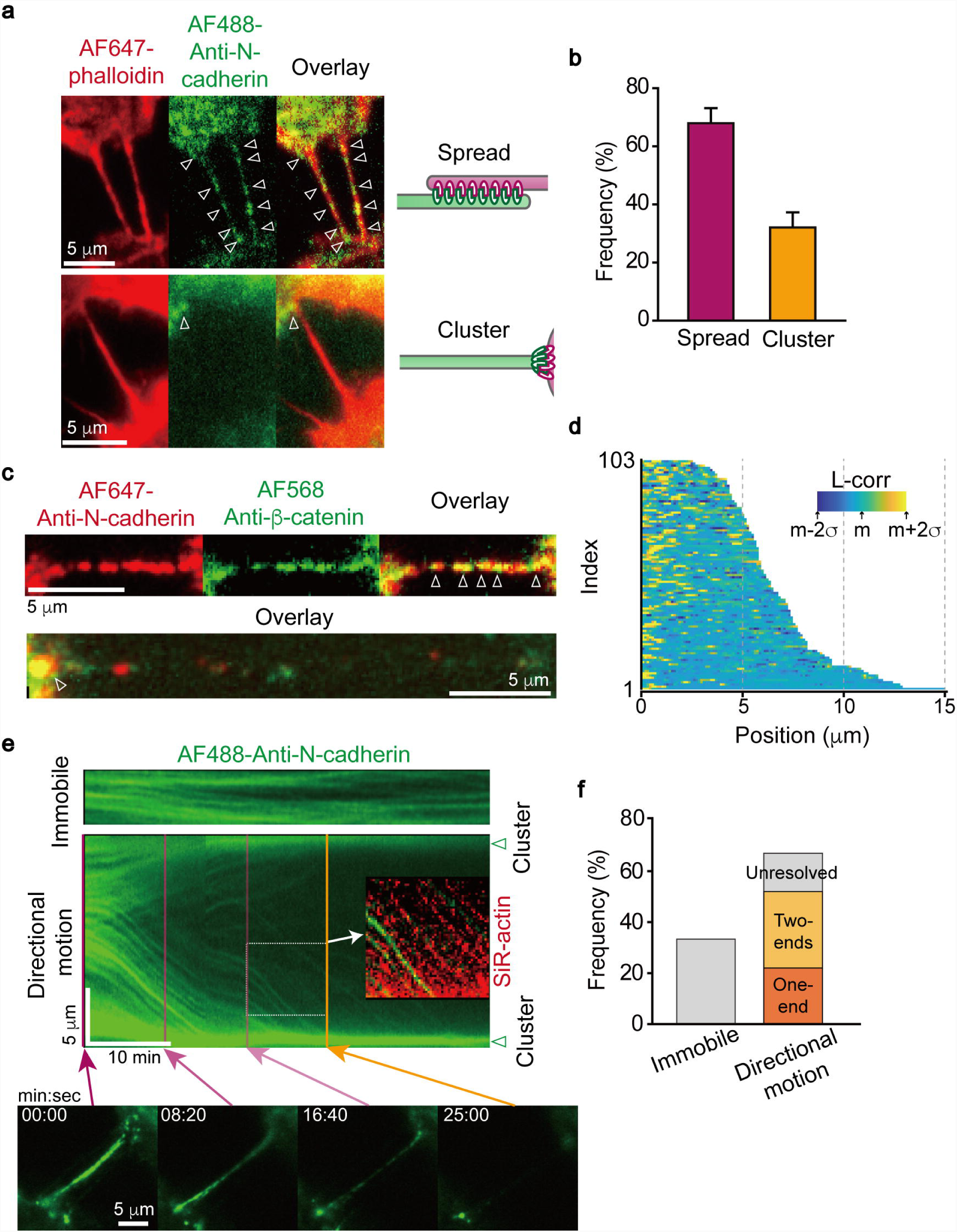
Cadherin-cadherin interactions form FBs and connect INTs to cells. **a**, Colocalization of N-cadherin (anti-N-cadherin antibody conjugated to AF488) and F-actin (AF647-phalloidin) shows that N-cadherin molecules are randomly distributed along the intercellular bridge between cells (Spread) or clustered at the junction of the bridge and the paired cell body (Cluster). **b**, The frequency of the N-cadherin distribution on FBs or INTs between cells (n = 143, N_cell_ = 124, N_exp_ = 4). The cells were fixed after 12 hr in stimulating conditions 24 hr after cell seeding. The error bars indicate the s.d. **c**, N-cadherin/β-catenin complexes in the fixed cells are visualized using an anti-N-cadherin antibody conjugated to AF647 and an anti-β-catenin antibody conjugated to AF568. **d**, The local correlation distribution of N-cadherin/β-catenin complexes on each FB or INT (n = 103, N_cell_ = 67, N_exp_ = 2). The colour code represents the strength of the local correlation at each pixel [L_corr(i), Methods; mean = 0.010, s.d.= 0.026 of all the pixels of an FB or INT]. We divided the FB or INT in half, compared their local correlations, and then aligned the FB or INT such that the half with the stronger correlation started from the origin. **e,** Kymographs of N-cadherin motion based on real-time images representing the immobile state or directional motion on FBs or INTs in living cells (n = 27, N_cell_ = 19). Actin filaments were labelled with SiR-actin for live cell imaging. The interaction of N-cadherin/β-catenin complexes bound to each filopodium forming the FB may result in an immobile state (top panel). A significant number of N-cadherin molecules move to the cell periphery (middle and bottom panel). The directional motion of N-cadherin was synchronized with F-actin retrograde flow (Inset). **f**, The frequency of N-cadherin in the immobile state (33%) and exhibiting directional motion (67%) (n = 27, N_cell_ = 19). Among the N-cadherin molecules exhibiting directional motion, an N-cadherin cluster appeared to be at an end (22 %) or at both ends (30%). Unfortunately, clusters were not resolved in some cells due to the bright cell body (15%). The real-time images were obtained from a time-lapse experiment (recording every 5 or 10 sec with a 100 ms exposure time).

Real-time imaging in live cells showed the immobile or unidirectionally mobile state of N-cadherin molecules on the bridge between two cells (Fig. 2e; Supplementary Movie 4). The immobile N-cadherin molecules may result from strong mutual adhesion between filopodia in the FB. In contrast, the mobile N-cadherin molecules moved to the cell body and then appeared to accumulate at the junction of the end of the filopodium and the cell body (Fig. 2e, bottom). This directional motion was correlated with retrograde F-actin flow (Fig. 2e, Inset) and led to a cluster at one end (22%) or clusters at both ends (30%) (Fig. 2f). This result can be interpreted to indicate that the interaction of N-cadherin between filopodia, which forms a complex with β-catenin bound to F-actin, was broken during the transition of FBs. We carried out a Ca^2+^ depletion experiment with 2 mM EGTA for 30 min or 20 mM EGTA for 3 min to interrupt the interaction between the extracellular domains of N-cadherin molecules (Supplementary Fig. 4)^42, 43^. The frequency of the filopodial separation in FBs was significantly enhanced after Ca^2+^ depletion, which provides strong evidence that FBs form via the dimerization of N-cadherin extracellular domains from different filopodia. Taken together with the observed stability of INTs bound to the other cell body even under Ca^2+^ depleted conditions (Supplementary Fig. 4c), these results suggest that N-cadherin/β-catenin clusters maintain the strong connection between INT and the cell body.

### Myosin II and V regulate INT formation via retractive force and torque

Surprisingly, we observed that FBs had a helical structure after visualization using real-time super-resolution microscopy (Fig. 3a; SRRF-stream, Methods). The rotational retraction of one filopodium in an FB is likely to result from the twisting of one filopodium while the other filopodium remains connected to the paired cell body (Supplementary Fig. 5; Supplementary Movie 5).

**Fig 3.**
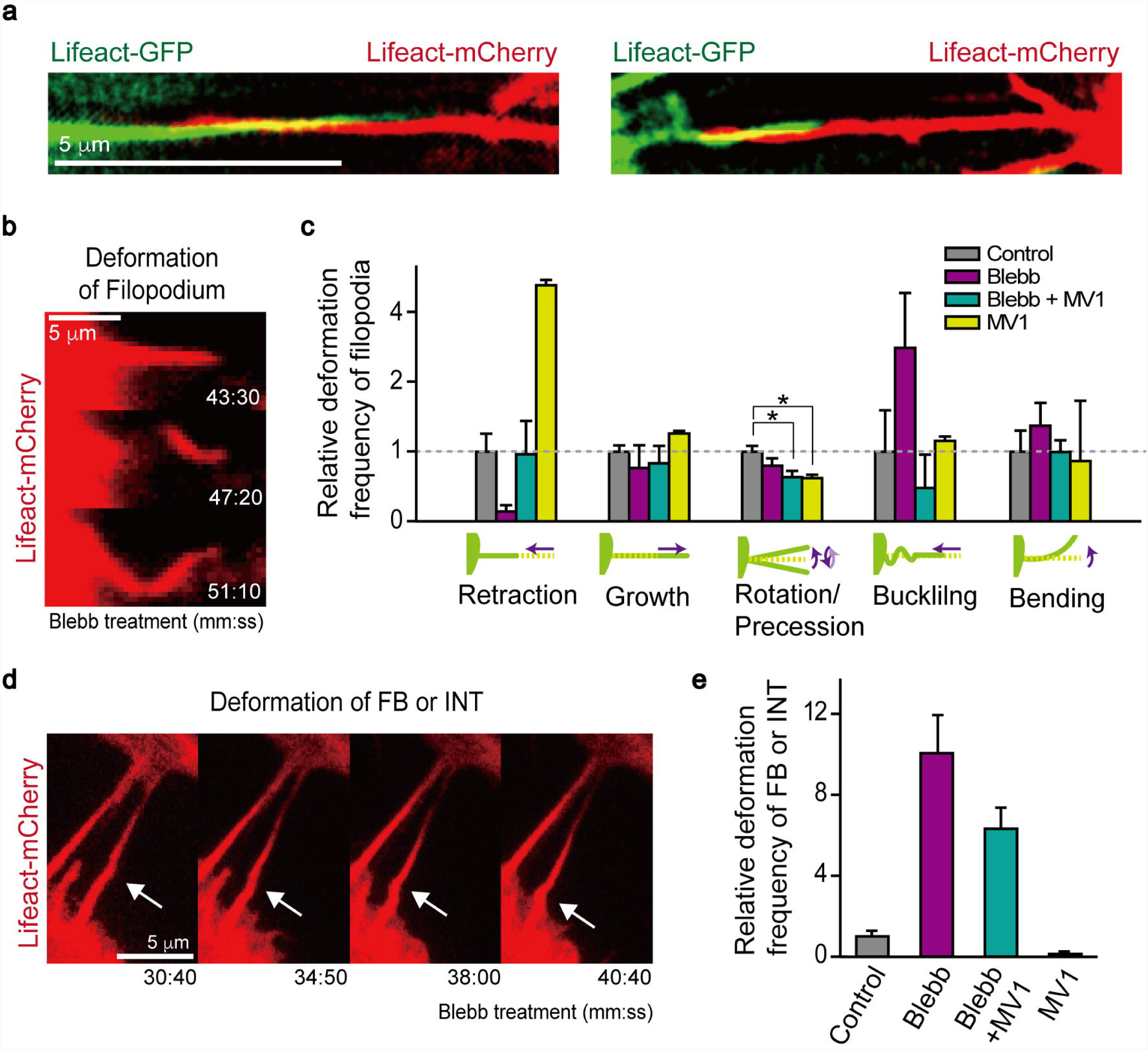
Myosin II and V regulate the FB helical structure. **a**, Twisted structures of FBs in living HeLa cells that were imaged by SRRF microscopy (iXon SRRF-Stream, Andor). Lifeact-GFP or - mCherry was transfected into HeLa cells for F-actin staining. Images were taken after 12 hr of INT stimulation 12 hr after cell seeding. **b**, Myosin II inhibition by 100 μM (±)-blebbistatin (Blebb) results in helical buckling and a precession (rotational) motion of filopodia. **c**, Relative deformation frequency of filopodia in the presence of Blebb (n = 100, N_exp_ = 4), Blebb + MV1 (n = 77, N_exp_ = 3), or MV1 (n = 55, N_exp_ = 2) compared to the deformation of filopodia without the addition of any drug (control, n = 55). Filopodial motion prior to any drug treatment was described as retraction, growth, precession/rotation, helical buckling, or bending. After a 30-min treatment with Blebb or MyoVin-1 (MV1) at the same concentrations to inhibit myosin V, myosin II inhibition reduces the retraction and significantly increases helical buckling, while myosin V inhibition dramatically increases retraction and reduces precession/rotation. \**p*-value < 0.05 by Student’s t-test. The error bars indicate the standard error (s.e.). **d**, Buckling of FBs or INTs after Blebb treatment. **e**, Relative deformation frequency of FBs or INTs after treatment with Blebb (n = 165, N_exp_ = 4), Blebb + MVI (n = 167, N_exp_ = 3), or MVI (n = 220, N_exp_ = 3) compared to the deformation of FBs or INTs in the absence of any drug (control, n = 139, N_exp_ = 2). The error bars indicate the s.e.

F-actin dynamics can drive various conformations of filopodia, such as retraction, growth, rotation/precession, helical buckling, and bending (Fig. 3b). Treatment with (±)-blebbistatin (Blebb), which inhibits myosin II, dramatically suppressed the retraction of filopodia (∼10-fold), but MyoVin-1^44^ (MV1), which inhibits myosin V, enhanced the retraction of filopodia (∼3-fold) (Fig. 3c). These observations are consistent with the previous results^45^. The precession/rotation conformation was observed ∼2-fold less frequently after MV1 treatment than in the untreated cells (Fig. 3c). The frequency of helical buckling, which can occur due to the axial rotation of F-actin inside the filopodium^36^, increased up to ∼ 2.5-fold when myosin II was inhibited (Supplementary Movie 6), but decreased ∼ 2-fold after the inhibition of both myosin II and V (Fig. 3c). These results indicate that myosin II reduces the helical buckling of F-actin by pulling the actin molecules into the cells, while myosin V promotes helical buckling by inducing axial rotation.

Remarkably, we also observed conformational changes in FBs and INTs after treatment with Blebb and/or MV1. Fig. 3d shows real-time images of rotation-coupled conformations of FBs (or INTs), such as helical buckling and precession/rotation, after the inhibition of myosin II (Supplementary Movie 7). The drug treatments did not change the frequency or length of FBs or INTs (Supplementary Figs. 6a, b). Blebb and Blebb/MV1 increased the rotation-coupled conformations by ten- and six-fold, respectively, while MV1 decreased the rotation-coupled conformations by 10-fold (Fig. 3e). Taken together, these observations suggest that the dynamics of FBs are associated with non-muscle myosin II, which exerts a retractive force on the F-actin inside the filopodium, and myosin V, which induces right-handed rotations in the F-actin in the filopodium^35^.

## Helical twisting of FBs promotes the transition to INTs

The nanoscaled structure of FBs and INTs remains unresolved due to the poor spatial resolution of conventional fluorescence microscopy (Fig. 4a, top). We employed direct stochastic optical reconstruction microscopy (dSTORM; Methods) to visualize the nanostructure of FBs and INTs in fixed HeLa cells. The dSTORM images showed the helically twisted filopodia of FBs as shown in Fig. 3a and 4a, and provided the finer structure, while INTs are visualized as linear bridges connecting two cells (Fig. 4a, bottom). We confirmed that the helical structure was not an artefact due to phalloidin labelling by showing the identical helical structure of FBs in silicon rhodamine (SiR)-actin-stained cells^46^ (Supplementary Fig. 7). To analyse the helical structure of FBs, we determined the peak position of the double Gaussian fitting of the AF647-phalloidin intensity profile in fixed cells (Fig. 4b). The minimum separation between filopodia was resolved based on the 20% contrast of the Rayleigh criterion (Fig. 4b, cartoon)^47^. The helical pitch and the distance between filopodia in FBs, which varied periodically along the lateral surface of the FB, were measured as 1,201 ± 994 nm (mean ± s.d.) and 232 ± 64 nm (mean ± s.d.), respectively (Supplementary Fig. 8). In addition, the release of the twisted structure of FBs after MV1 treatment (Supplementary Fig. 9) indicates that the rotational motion of F-actin by myosin V is likely to contribute to the helical structure of FBs. Figs. 4c, d, and e show dSTORM images of FBs, FBs transitioning to INTs, and INTs. The N-cadherin molecules corresponding to each state appeared to be located at the end of the bridge as the FBs were transformed into INTs, as shown in Fig. 2e (Figs. 4c, d, e). The intensity of N-cadherin molecules at the end of the partial helical structure in contact with the cell body was stronger than that at the junction of the single protrusion and the other cell body (Figs. 4d, e, inset), which indicates that the filopodium weakly bound to the cell body was released. This super-resolution imaging reveals the physical characteristics of the helical structure of FBs and the nanostructure of the intermediate state during the transition of FBs to INTs.

**Fig 4.**
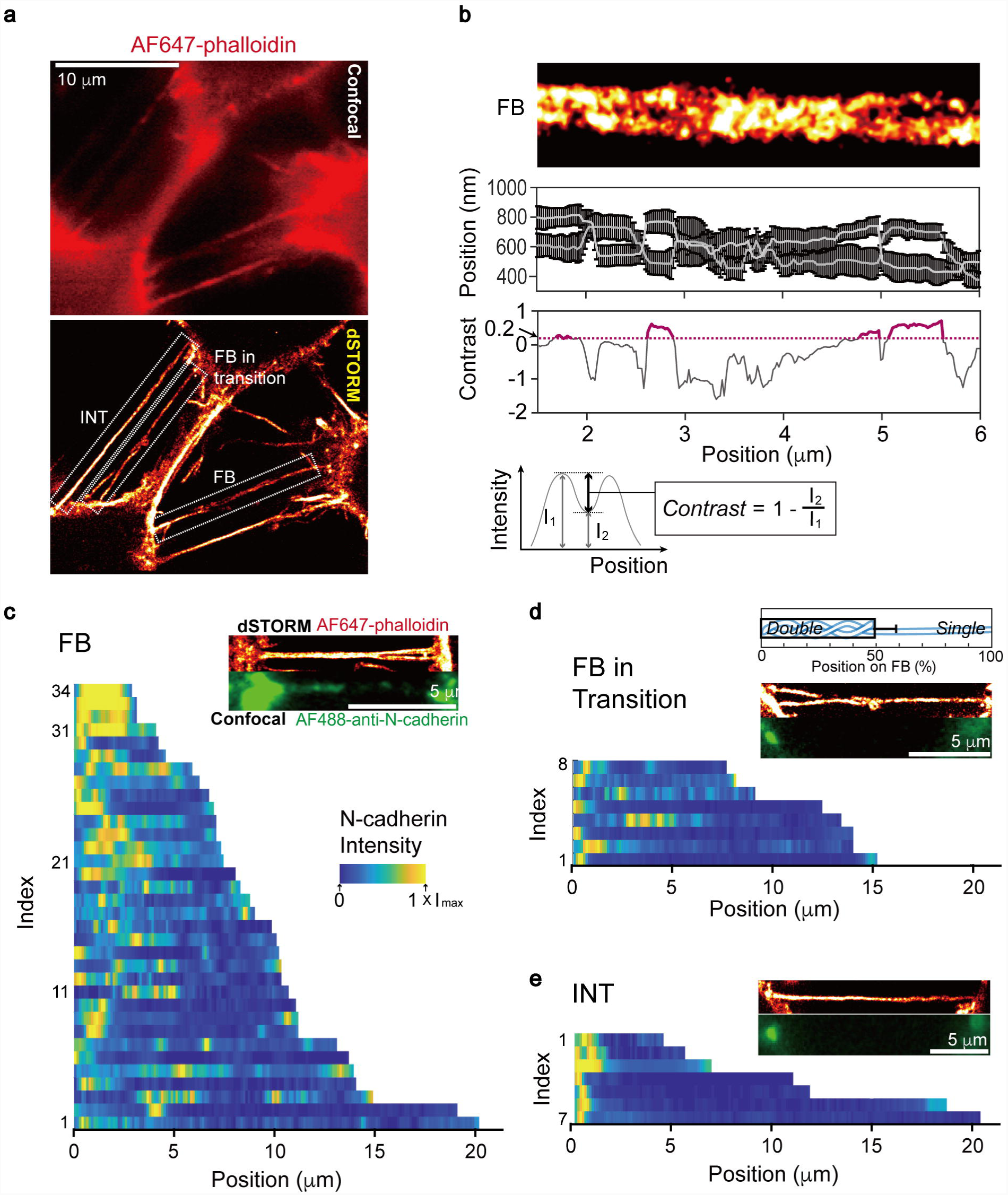
Super-resolved nanostructure of FBs and INTs. **a**, Super-resolution microscopy (dSTORM) reveals heterogeneous structures of FBs or INTs labelled with AF647-phalloidin in fixed cells (bottom panel), which were indistinguishable by confocal microscopy (top panel). We observed 77% FBs and 23% INTs among 115 connections obtained from 66 cell pairs. **b**, The centre position determined by a double Gaussian fitting of the intensity profiles of AF647 (middle panel) in a dSTORM image (top panel). The error bars indicate the s.d. (middle panel). The minimum resolvable space between filopodia forming an FB was determined by the 20% contrast of the Rayleigh criterion (bottom panel). **c, d, e,** FBs (c), FBs in transition to INTs (d), INTs (e), and N-cadherin distributions corresponding to the states of the connections. By using AF647-phalloidin (for dSTORM) and an AF488-conjugated anti-N-cadherin antibody (for confocal microscopy), fixed cells were imaged simultaneously. N-cadherin molecules were clustered at the junction of the INT and the acceptor cell body. The intensity distributions of N-cadherin molecules on FBs (n = 34; c), FBs in the transition to INTs (n = 8; d), and INTs (n = 7) (N_cell_ = 27; e). Each pixel intensity for an FB or INT was normalized based on the maximum pixel intensity of each profile and represented by the colour code (inset). The FBs in transition to INTs were partially twisted (49.2 ± 9.4% of the total length of the bridge between cells) on a single protrusion connecting cells.

## Discussion

Since INTs have been recognized as a possible conduit for long-distance communication between cells, there have been many efforts to understand the biological roles of INTs by studying the transport of cellular components at the cellular level and identifying the molecules that induce the formation of INTs. Nonetheless, the mechanisms that underlie the formation and maintenance of actin-driven INTs have not yet been revealed. Our real-time fluorescence and super-resolution imaging experiments revealed that the spatiotemporal processes of INT formation evolved from the fine nanostructure of FBs. Here, we propose a mechanical model for INT formation from the initial stage (random contact between two filopodia) to the functional configuration (a single bridge connecting two cells) through an intermediate state (the twisting of two filopodia) for the ultimate construction of a specific signaling structure.

The extracellular interaction between N-cadherin molecules on two distinct filopodia that are linked to β-catenin sustains the physical contact between the filopodia in FBs (Fig. 2a). However, filopodia often fail to make a stable connection due to fluctuations caused by the active motion of F-actin inside the filopodia (Supplementary Movie 1). The critical adhesion between two filopodia may be required for the formation of an FB and ultimately for the transformation to an INT. In fact, since the N-cadherin/β-catenin complex inhibits F-actin retrograde flow^48^, it can direct the growth of the filopodia in an FB. Interestingly, N-cadherin molecules moved to the end of filopodia in FBs, while INTs formed from FBs (Fig. 2e, Supplementary Movie 4), in which N-cadherin molecules are supposed to be transported in the anterograde direction. We suspect that myosin X, which is required for filopodium formation and extension, can deliver N-cadherin to the tip of the filopodium, as found in neuronal cells^49-51^, in contrast to the retraction of the filopodium during the transition of FBs. Since we did not observe the significant breakage of INTs once they were connected to the paired cell body, even under Ca^2+^ depleted conditions (Supplementary Fig. 4), we concluded that the N-cadherin/β-catenin cluster at the filopodium end provides a strong bond between the end of the filopodium and the paired cell body.

How is one of two filopodia then released from the other filopodium or the other cell body to form a single protrusion bridge (INT)? The unexpected finding in our study is the observed twisting of filopodia in FBs (Fig. 3a: Fig. 4; Supplementary Fig. 10), which results from the rotation of F-actin inside the filopodium by myosin V (Fig. 3; Supplementary Fig. 9). Since the distance (232 nm) between the antinodes of the helical FB is greater than the thickness of a single filopodium (188 nm) on average, the N-cadherin interactions between two filopodia are likely to be broken at the antinodes (Supplementary Fig. 8). We speculate that the torsional energy that accumulates in the helical FB can locally separate two filopodia and even one filopodium weakly bound to the cell body. The filopodium released from the cell body or the other filopodium is then retracted by the F-actin retrograde flow inside the filopodium (Fig. 1b; Fig. 4c; Supplementary Fig. 5).

We observed vesicles moving to the paired cell body along INTs in membrane-labelled cells^9^ (Supplementary Figs. 11a, b) and epidermal growth factor receptors (EGFR) molecules unidirectionally moving on INTs in our previous study^52^. The rate of the EGFR transport was decreased globally when an external force was applied in the opposite direction of the transport, which suggests that EGFR transport on INTs resulted from the retrograde flow of F-actin, as reported for filopodia^53^. In fact, unidirectional actin flow away from the cluster is visible on INTs (Supplementary Fig. 11c), which explains why vesicles and EGFR molecules were transported by the F-actin retrograde flow along INTs. The cellular components in a donor cell are transferred to an acceptor cell along INTs via a specific junction (INT synapse) at the site of contact between the INT and a cell body^54-57^ or an open end of the INT (tunnelling nanotube)^6^. Although we could not reveal the fine structure of the INT terminus that physically contacts a cell body through N-cadherin molecules^58^, we speculate that the N-cadherin cluster plays an important role in forming INT synapses between the end of the INT and the cell body since N-cadherin molecules are often found at the synapse in neuronal and immune cells^59, 60^.

In summary, we propose a mechanical model for INT formation by actin-based protrusions in HeLa cells (Fig. 5). (1) Two filopodia that protrude from the cells involved in cell-to-cell communication appear to make physical contact through N-cadherin molecules that are anchored to F-actin via catenin molecules. (2) A high density of N-cadherin/catenin complexes inhibits retrograde F-actin flow (solid arrow). Instead, the filopodia are extended by actin polymerization until the filopodium reaches the paired cell body. (pointed arrow = anterograde transport by myosin X). (3) The torsional energy given to F-actin by myosin V results in the helical structure of the FB. The accumulated torsional energy forces the two filopodia to separate, which allows the transportation of N-cadherin molecules to the junction of the filopodium and the cell body. (4) One filopodium is released and retracted. (5) The cluster of N-cadherin/catenin complexes holds the INT at the junction and forms a synapse between the INT and the cell body for distant communication between cells.

**Fig 5.**
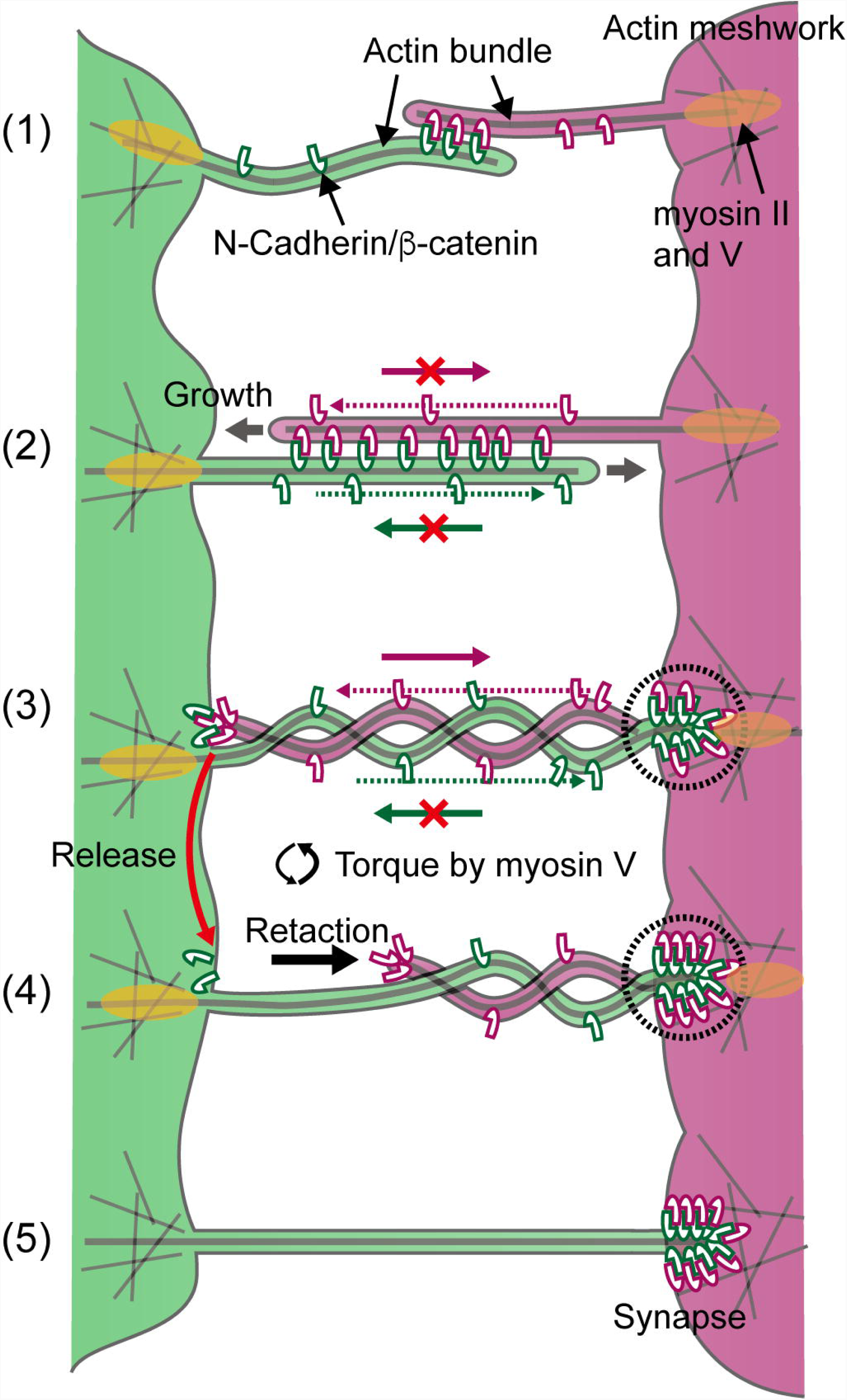
A mechanical model of INT formation by actin-based protrusions. See the text in the Discussion for details.

## Acknowledgements

This research was supported by the National Research Foundation of Korea funded by the Ministry of Science and ICT (2017R1A2B3006006 to J.D and 2017K1A1A2013241 to J.-B.L).

## Author contributions

M.C. and J-B.L design the experiments; M.C. performed all the experiments except dSTORM imaging: J.O. performed the dSTORM imaging; J.D. provided F-actin labelling materials; M.C., J.O., J.D. and J.-B.L. analyzed the data; All authors participated in critical discussions. M.C. and J.-B.L. wrote the paper.

## Additional information

Supplementary information is available for this paper at….

## Methods

### Fluorescence imaging

Video-rate LSCM (Supplementary Fig. 1) was used for fluorescence imaging, as previously described^52^. We collected the emissions from excited fluorophores through a 60× water-immersion (Olympus UPlanSApo 60×, N.A. = 1.2, for the formation-transition study, Fig. 1) or 100× oil-immersion objective (Olympus UPlanSApo 100×, N.A = 1.4, for the other studies) on an inverted microscope (Olympus IX51). For multi-colour excitation, up to 4 different lasers (Cobolt MLD^TM^ 405 nm, 100 mW; Cobolt MLD^TM^ 488 nm, 60 mW; CNI DPSS laser 561 nm, 200 mW; Cobolt MLD^TM^ 638 nm, 100 mW) were introduced sequentially (for the formation-transition study and time-lapse imaging of N-cadherin in Figs. 1, 2e) using mechanical shutters (Uniblitz, LS3S2T0 and VMM-D3) or simultaneously (for simultaneous imaging of N-cadherin and β-catenin in Fig. 2c or N-cadherin and F-actin via dSTORM in Figs. 2a, 4c). Different excitation and emission wavelengths were reflected and transmitted, respectively, through a quad-edge dichroic beam splitter (Semrock, Di01-R405/488/561/635). Galvano scanning mirrors (Cambridge Technology 6231H, 15 mm) were controlled using a homebuilt control program (LabView). The emission signals were separated temporally through a multiple bandpass filter (Semrock, FF01-515/588/700-25) (for sequential multi-excitations) or spatially through a DV2 multichannel imaging system (Photometrics) with suitable bandpass filters (Semrock, 510/20 for AF488, 600/37 for AF568, 680/40 for AF647 and SiR-actin) (for simultaneous multi-excitations) in front of an electron multiplying charge-coupled device (EMCCD) camera (Andor iXon or iXon Ultra). Separated signals were obtained using imaging software (Solis or MetaMorph) with a time resolution of 100 ms. For the time-lapse experiment using sequential multi-excitations, the time interval was mainly set as 9.9 s for 60-180 min. To maintain cell viability during live cell imaging, a heated sample stage and a CO_2_ supply system (Live Cell Instrument) were used. The image sequences obtained from the imaging program were analysed using ImageJ and MATLAB (MathWorks) software. To overlay multi-colour images of simultaneous multi-excitations, fluorescent bead images were recorded as a reference for mapping. Images were occasionally walking-averaged over every four frames. For SRRF microscopy (Fig. 3a), we used the imaging method for the iXon SRRF-Stream (Andor).

### Actin and membrane imaging

For F-actin labelling, 1 μg of the Lifeact-GFP or Lifeact-mCherry plasmid^38^ (Addgene) was transfected into HeLa cells and incubated for 24 hr after subculturing (80∼90% confluence) using 2 μl of the Lipofectamine 3000 reagent (Thermo Fisher Scientific) on 35 mm dishes. The cells were cultured in phenol-red free DMEM to reduce autofluorescence (10% FBS, 1× Pen Strep, 1× Glutamax). The transfection medium was changed after 12 hr of transfection. The transfected cells on both dishes were trypsinized 12 hr after the media change and gathered into a 1.7 ml micro-centrifuge tube for gentle centrifugation (2 min, 1500 g). After the supernatants were removed, the cells were suspended in different volumes of medium in a poly(dimethylsiloxane) (PDMS) chamber for live cell imaging^52^ or a 4-chamber glass bottom dish (Cellvis, fixed cell or live cell imaging) to control the cell density. We mixed the cells by repeatedly dispensing them using 18G syringe needles and then injected the mixed-cells, generally 400 μl, into a PDMS chamber or a glass-bottom dish for imaging. For membrane labelling, DiI and DiD (Molecular Probes) were diluted to 5 μM in medium and used to treat cells on 35 mm dishes (24 hr after subculture) separately for 8 min and then washed out according to the manufacturer’s protocol. The cells were immediately mixed and moved to a PDMS chamber or a glass-bottom dish for imaging. The cells were imaged 36 to 60 hr after mixing and seeding. The medium was replaced with INT formation-stimulating medium (2.5% FBS in DMEM, 50 mM glucose, pH 6.6)^23^ 12-24 hr after cell seeding. For fixed cell imaging, the cells were fixed with 4% paraformaldehyde in cytoskeleton-preserving buffer (80 mM PIPES pH 6.8, 5 mM EGTA, 2 mM MgCl_2_) for 10 min, washed with PBS and contained in PBS for imaging. We used (±)-blebbistatin (Calbiochem and Abcam) and MyoVin-1 (Calbiochem) to inhibit myosin II and V, respectively. The cells were subjected to F-actin-labelling after 12-24 hr of INT stimulation. For live cell imaging, the cells were treated with a 100 μM concentration of one or both drugs in medium. For fixed cell imaging, the cells were fixed after 30 min of drug treatment.

### N-cadherin and β-catenin imaging

N-cadherin was labelled with an anti-N-cadherin antibody conjugated to AF488 or AF647 (Clone 8C11, BD). Two microlitres of AF488- or AF647-conjugated anti-N-cadherin antibody was diluted with 300 μl of medium. For living cells, actin filaments were incubated with 200 nM SiR-actin (Spirochrome) for 3 hr at 37^°^C and AF488- or AF647-conjugated anti-N-cadherin antibody and SiR-actin were then simultaneously injected and incubated in the imaging chamber for 30 min at 37^°^C. After washing twice with pre-warmed PBS, the cells were imaged with 200 nM SiR-actin to maintain F-actin labelling during time-lapse imaging. For fixed cell imaging, after the cells were incubated with an AF488- or AF647-conjugated anti-N-cadherin antibody and an AF568-conjugated anti-β-catenin antibody (E247, Abcam) for 30 min at 37^°^C, and the unbound antibodies were washed out. The cells were fixed with 4% paraformaldehyde in cytoskeleton-preserving buffer. The fixed cells were rinsed with PBS and then permeabilized with 0.5% Triton-X-100 (Sigma) in PBS for 10 min after a 0.5 ∼ 1 hr blocking step with 5% (w/v) bovine serum albumin (BSA; Sigma) at room temperature. For actin filament labelling, the fixed cells were incubated in a 200-400 nM AF647-phalloidin (Thermo Fisher Scientific) solution overnight at 4°C.

### Direct STORM imaging

Direct STORM (dSTORM) imaging was performed under LSCM with an objective (Olympus, UPlanSApo 100×, N.A. = 1.4, oil immersion) that results in a 120 × 120 nm pixel size (Supplementary Fig. 1). Actin filaments labelled with AF647-phalloidin or SiR-actin were imaged with a 10-20 Hz video rate for 5,000-20,000 frames. We used weak excitation (0.04 ∼ 0.1 kW/cm^2^) with a 638 nm DPSS laser to visualize FBs or INTs. For dSTORM imaging, we increased the excitation power (∼3.5 kW/cm^2^) to rapidly activate the fluorophore and enhance the positional accuracy with the number of emitted photons. A 405 nm laser was used to reactivate the fluorophore in the dark state with a weak excitation power (0 ∼ 0.35 kW/cm^2^). The acquired images were analysed with ImageJ and ThunderSTORM^61^ to localize the switching spots. The analysed localization data were rendered with the normalized Gaussian option in ThunderSTORM with a 12 × 12 nm pixel size to accurately determine the position of each spot. The background noise was filtered for the optimal imaging condition, and the X-Y drift was corrected using the cross-correlation function in ThunderSTORM. The images were coloured with ImageJ Red Hot. The intensity profile of a single AF647 molecule has a full width at half maximum (FWHM) of ∼ 23 nm. The cells were fixed with 4% paraformaldehyde in cytoskeleton-preserving buffer and incubated with a fresh 0.1% (w/v) sodium borohydride (Sigma-Aldrich) solution for 7 min to reduce aldehyde-related fluorescence. After a 0.5 ∼ 1 hr blocking step with 5% (w/v) BSA (Sigma) at room temperature, the cells were incubated with a 200-400 nM AF647-phalloidin (Thermo Fisher Scientific) solution overnight at 4^°^C for actin filament labelling. The labelled samples were rinsed once with PBS. The solution was changed to imaging buffer [100-200 mM β-mercaptoethanol, 2 mM Trolox, 2.5 mM protocatechuic acid (PCA) and 50 nM protocatechuate-3,4-dioxygenase (PCD) in PBS]. For the SiR-actin probe experiment, cells grown in a PDMS chamber for 24 hr were fixed and labelled with SiR-actin probes (250 nM ∼ 750 nM) for 1 hr after the Triton X-100 treatment (10 min) and BSA pre-incubation (5% BSA/PBS). After the free probes were washed out (PBS, 0.1% Tween20), imaging was carried out in the presence of imaging buffer. The intensity profile of a single SiR has an FWHM of ∼ 33 nm.

### Local correlation of N-cadherin and β-catenin

We measured the intensities of AF647-N-cadherin (INcad) and β-catenin (*I*_cat_) on the FBs or INTs between cells, for which the background intensity was subtracted from the colocalization image. The global correlation (Corr) was measured by Pearson’s correlation coefficient

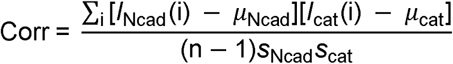

for the mean (*μ*) and the standard deviation (*s*) of each profile. The local correlation at each pixel *i* (L_corr(i)) along an FB or INT was measured by

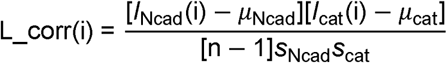

where the mean (*μ*), the standard deviation (*s*) of each profile, and the total number of pixels for the FB or INT were used.

